# Atypical DNA methylation, sRNA size distribution and female gametogenesis correlate with genome compaction in *Utricularia gibba*

**DOI:** 10.1101/2020.12.05.413054

**Authors:** Sergio Alan Cervantes-Pérez, Lenin Yong-Villalobos, Nathalia M. V. Florez-Zapata, Araceli Oropeza-Aburto, Félix Rico-Reséndiz, Itzel Amasende-Morales, Tianying Lan, Octavio Martínez, Jean Philippe Vielle-Calzada, Victor A. Albert, Luis Herrera-Estrella

## Abstract

- The most studied DNA methylation pathway in plants is the RNA Directed DNA Methylation (RdDM), which is a conserved mechanism that involves noncoding-RNAs to control the expansion of intergenic regions. However, little is known about relationship between plant genome size reductions and DNA methylation.
- Because the compact genome size of the carnivorous plant *Utricularia gibba,* we investigate in this plant the noncoding-RNA landscape and DNA methylation through a combination of cytological, evolutionary, and genome-wide transcriptomic and methylation approaches.
- We report an unusual distribution of noncoding RNAs in *U. gibba* in comparison with other characterized angiosperms, which correlated with a lower level of global genome methylation, as determined by a novel strategy based on long-read DNA sequencing and corroborated by whole-genome bisulfite analysis. Moreover, found that genes involved in the RdDM pathway may not be functionally active in *U. gibba*, including a truncated *DICER-LIKE 3 (DCL3),* involved in the production of 24-nt small-RNAs.
- Our findings suggest that selective pressure to conserve a fully functional RdDM pathway might be reduced in compact genomes and a defective *DCL3* correlate with a decreased proportion of 24-nt small-RNAs and developmental alterations in *U. gibba*, which could represent an initial step in the evolution of apomixis.

## Introduction

Epigenetic modifications are chemical additions to DNA and/or histones that are associated with changes in gene expression and can be heritable (Law & Jacobsen, 2010). One of these epigenetic modifications is DNA methylation at the 5ʹ position of the cytosine base (m5C), an ancient evolutionary trait associated with gene and transposable element (TE) silencing in eukaryotes (Slotkin & Martienssen, 2007). In plant genomes, m5C is a widely conserved epigenetic mark that modulates gene expression and plays a key role in many developmental processes and environmental responses (Henderson & Jacobsen, 2007; Zhang *et al.*, 2018).

Non-coding RNAs (ncRNAs) are fundamental in regulating DNA methylation and the accessibility to genetic information (Wierzbicki *et al.*, 2008). For instance, mobile short RNAs (sRNAs) underlie shoot to root communication of methylation status (Lewsey *et al.*, 2016) and long-noncoding RNAs (lncRNAs) serve as scaffolds in *de novo* DNA methylation (Ariel *et al.*, 2015). RNA-directed DNA methylation (RdDM) is the major small RNA-mediated epigenetic pathway in plants. The canonical pathway can be subdivided into 3 different phases: 1) RNA polymerase IV (Pol-IV) dependent biogenesis of small interfering RNAs (siRNAs), 2) RNA polymerase V (Pol-V) mediated *de novo* methylation, and 3) chromatin modifications (Matzke & Mosher, 2014). In addition to Pol-IV and Pol-V, other key components in RdDM and sRNA biogenesis pathways are the ARGONAUTE (AGO), DICER-LIKE (DCL), and RNA DEPENDENT RNA POLYMERASE (RDR) gene families. Specifically, in the production of siRNAs, RDRs synthesize the second RNA strand, DCLs process RNA precursors, and AGOs select one DNA strand and load sRNAs to a specific target (Bologna & Voinnet, 2014; Matzke & Mosher, 2014). A considerable number of plant methylomes have been analyzed by whole-genome bisulfite sequencing (BS-Seq) or by high-performance liquid chromatography (Alonso *et al.*, 2015; Vidalis *et al.*, 2016), of which the vast majority of these methylomes are from model flowering plants. Methylome analysis has revealed variation of methylation patterns in intergenic and gene body regions among different plant species (Niederhuth *et al.*, 2016; Vidalis *et al.*, 2016) and these patterns could be related to genome architectural features such as genome size, rearrangements, duplications, and content/expansion of transposable elements (TEs), among others. Little is known about the relationship between genome methylation, number, type of ncRNA *loci*, and genome size, particularly for plant species with small genomes with reduced content of TEs and other repetitive sequences.

*Utricularia gibba* is an aquatic, carnivorous plant belonging to the asterid lineage that despite having undergone two recent whole genome duplication (WGD) events, has a remarkably small genome size (ca. 100 Mb) (Ibarra-Laclette *et al.*, 2013). *U. gibba* is a rootless plant that harbors slender green stems that grow like stolons with alternate thread-like leaves and numerous complex trapping bladders that catch small invertebrate prey. *U. gibba* has a gene repertoire similar to other plants species with larger genomes, albeit with a reduced non-coding genome harboring short intergenic regions (IGR) and a low TE content (Ibarra-Laclette *et al.*, 2013). We took advantage of the small *U. gibba* genome to study the impact of genome size on the conservation of “non-coding DNA”, especially on nature of genes coding for ncRNAs and the impact of the ncRNA landscape on genome methylation. Here, we report that *U. gibba* has, compared to Arabidopsis, a functionally incomplete canonical pathway for siRNAs biogenesis and *de novo* methylation that correlates with a rather unusual content and proportion of sRNA. In addition, our data suggests the interesting hypothesis that the unusual siRNA content and altered female gametogenesis in *U. gibba* could be due to the lack of a functional *DCL3*. Finally, we proposed that Pacific Bioscience (PacBio) Single Molecule Real-Time (SMRT) long read sequencing data can be used to estimate the content and distribution of m5C methylomes in plant genomes.

## Material and Methods

### Plant material

As in previous studies, *U. gibba* was obtained from Umécuaro village in the municipality of Morelia, México. Collected plants were grown in sterile tissue culture with media MS (0.25X) at 22 C with 16 hours light and 8 hours of darkness. Subcultures, at day 14 of growth, were used in our experiments. We employed the following growth conditions and treatments: nutrient deficiency, exposure to plant hormones, high and low temperatures, continuous illumination and darkness, treatment with acidic and alkaline pH, osmotic stress, and saline stress. Traps and plant tissue were collected separately after short (48 and 72 hours) and long (7 and 14 days) time treatments for total RNA extraction and subjected to RNA-Seq (Dataset S1).

### Total RNA extraction and sequencing

Total RNA from three independent biological replicates for all the analyzed treatments was isolated using the TRIzol reagent (Life technologies), except for trap tissue, which was isolated using the Direct-zol RNA kit (Zymo Research) because this protocol optimizes RNA extraction for low tissue quantities and improved RNA quality for these samples. Twelve RNA-Seq libraries were prepared with the TrueSeq (Illumina technologies) kit and sequenced by non-directional single-end mode.

### Mapping reads and lncRNAs identification

Trimmed reads were mapped against the latest available version of the *U. gibba* genome (https://genomevolution.org/coge/GenomeInfo.pl?gid=28800). To potentiate the identification of lncRNAs in *U. gibba* we decided to merge mapped reads for each library in a single file to obtain a core transcriptome assembly with Trinity v2.0 (Haas *et al.*, 2013). The identification works with three major filters: I) One of the main features for lncRNAs is size, then during the assembly the transcripts with less than 200 nucleotides in size were cut-off. The first step was to search protein orthologs against a database of non-redundant eukaryotic proteins (Suzek *et al.*, 2015), this step eliminates those transcripts that encode for proteins. II) Putative lncRNAs transcripts were filtered finding homologs with housekeeping RNAs such are transfer-RNAs (tRNAs), ribosomal-RNAs (rRNAs), micro-RNAs (miRNAs) and other types of structural RNA using the database Rfam v12.0 (Griffiths-Jones *et al.*, 2003) and sequence homologs with precursors of small RNAs from *U. gibba* dataset. III) Third step relays in the evaluation of the coding potential of the remaining transcripts using the tools CPC2 (Kang *et al.*, 2017) and CPAT v1.2.4 (Wang *et al.*, 2013) using the corrected training for *U. gibba* model. The identification is described and summarized in Fig. S1.

### Small RNA sequencing and annotation

RNA was extracted independently from each sample and then equimolarly pooled to produce three small RNA libraries (Dataset S1). Small RNA libraries were constructed using the TrueSeq Small RNA kit (Illumina Technologies) and sequenced using NextSeq (Illumina Technologies) using a 50-nucleotide single end runs. Cleaned reads were mapped against the *U. gibba* reference genome and the alignments were then run in ShortStack V2.0 (Axtell, 2013). ShortStack-count mode was used to find relative small RNA abundances of *de novo*-identified sRNAs *loci*. The sRNAs reads were mapped against mature sequences in the miRNA database miRBase V22.0 (blastn – word_size 7 –dust no –identity 90 –evalue 1000).

### Phylogenetic analysis

Were identified the homologs among plant genomes in Phytozome version 12.1 (http://phytozome.jgi.doe.gov/; (Goodstein *et al.*, 2012)). For all sequences, homology analysis was performed (Blastx E-value < 0.001, Bit-Score > 70). Additionally, we performed a bidirectional blast analysis against Arabidopsis protein database from TAIR V10 (www.arabidopsis.org) using previous parameters. Protein multiple alignments were made using MAFFT V7.0 (Katoh & Standley, 2013) and trimmed by trimAL (Capella-Gutiérrez *et al.*, 2009). Finally, the best model fit was selected to execute the phylogenetic inference with ProtTest V3.0 (Darriba *et al.*, 2011). The Bayesian phylogenetic reconstruction was executed using the software MrBayes V3.2 (Ronquist *et al.*, 2012) with 300000 generations. Also, likelihood phylogenetic reconstruction was estimated using RAxML back box on website CIPRES (https://www.phylo.org/) with GTR + G model and 1000 cycles for bootstrapping.

### Cytological analysis of ovule development

For cytological examination of ovules, whole flowers were harvested and fixed in formalin acetic acid-alcohol solution (40% formaldehyde, glacial acetic acid, 50% ethanol; in a 5:5:90 volume ratio) for 24 hours at room temperature, and subsequently stored in 70% ethanol at 4° C. Fixed ovaries were dissected with hypodermic needles (1 mm insulin syringes), cleared, and observed by differential interference contrast microscopy using a Leica DMR microscope.

### Identification of m5C with SMRT-sequencing and validation

Raw SMRT data was aligned against the reference genome using pbalign in base modification mode. Polymerase kinetics information was further loaded using loadPulses scripts and data was analyzed using the program SMRT-link V4.0 to identify base modifications. We used a filter of 15X coverage to select for m5Cs. The theoretical IPD value for non-modified bases is 1 (See Methods S1-S3). To validate our results, two genomic regions were selected from an entire chromosome, Unitig_0. Vegetative tissue explants were grown for 14 days and were transferred to new media with or without 50 μM of Zebularine for 8 days. DNA from this experiment was collected. The DNA was either digested or not with the methylation sensitive restriction enzymes Hae III or Hpa II. A quantification was performed using Chop-qPCR (Fig. S2).

### Whole-Genome Bisulfite Sequencing

Genomic DNA from *U. gibba* fresh tissue 14-days old was grinded with liquid nitrogen and purified with DNeasy Plant Maxi Kit (Qiagen). For the library preparation, DNA samples were fragmented into 200-400 bp using a sonicator S220 (Covaris). Then DNA fragments were blunt ended and, a dA 3′-end addition was performed prior to sequencing adapter ligation. Illumina methylated adapters were used according to the manufacturer’s instructions (Illumina). The DNA fragments were bisulfite treated with EZ DNA methylation Gold Kit (Zymo Research). The final DNA library was obtained by size selection and PCR amplification. High-throughput pair-end sequencing was carried out using the Illumina HiSeq 2500 system according to manufacturer instructions. The Clean reads were aligned to reference genome using Bismark software (Krueger & Andrews, 2011). To identify the true methylated sites, methylated and unmethylated counts at each site from Bismark output was tested by binomial distribution.

## Results

### *U. gibba* lncRNAs has a reduced number of intergenic lncRNAs

lncRNAs were obtained from sequencing data of 12 RNA-Seq libraries, 10 from vegetative and 2 from trap tissue from plants subjected to contrasting abiotic stress or hormone treatments (Dataset S1). We produced over 19 million mapped reads per library (95.37% mapping to the genome) for a total of 330,258,977 million mapped reads (Dataset S1). To identify lncRNAs, assembled transcripts were translated into the three potential reading frames and filtered by homology to proteins encoded in the *U. gibba* genome and other plants in a non-redundant protein database (Suzek *et al.*, 2015), yielding 10,386 putative lncRNAs of at least 200-nt in length. Putative lncRNAs were then filtered to eliminate precursors of tRNAs, rRNAs, miRNAs among others. Putative lncRNAs were further filtered for coding potential in sense-direction, resulting in 4,295 putative lncRNAs (Fig. S1). The vast majority of these lncRNA *loci* were relatively short, 89.05% were smaller than 500-nt, and only 1.81% longer than 800-nt (Fig. 1a). The lncRNA mean length was 336.9-nt and the largest one was 2134-nt (Table S1 and Dataset S2). We also found that in *U. gibba* 86.51% of lncRNAs had a single exon structure and 13.49% had two or more exons, percentages like other plant species for which lncRNAs have been reported (Fig. 1b; Dataset S1). When putative lncRNAs were mapped onto the *U. gibba* genome, we determined that 37.03% mapped to regions corresponding to exons of protein coding genes in antisense orientation, 10.89% to intronic regions, 25.26% overlapped gene bodies and IGRs, and 26.82% were located only in IGRs (Fig. 1c). As it is more complex to determine whether a non-coding transcript that completely overlaps with the transcript of a coding gene indeed corresponds to a lncRNA, we made only a comparison of intergenic lncRNAs (lincRNAs) with other plant species. By mapping directly RNA-Seq reads and assembled contigs, we found that this carnivorous plant, under all conditions tested, expresses a total of 1152 lincRNAs, which is significantly lower than that reported for other plant species that ranges between 1580 and 3100 lincRNAs (Fig. S3). Additionally, were evaluated the expression profiles for coding and noncoding transcripts and we found a higher level of expression for mRNA than for the entire catalog of lncRNAs or lincRNAs (Fig. 1d).

**Fig. 1.**
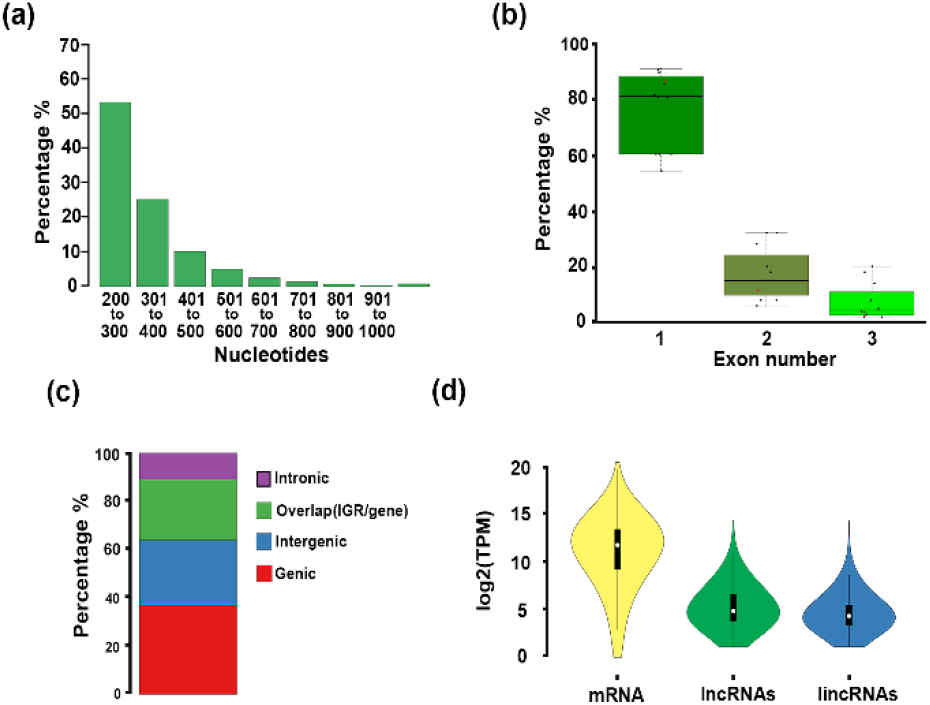
Architecture and annotation of lncRNAs in *U. gibba* and other plants. **(a)** lncRNAs size distribution in *U. gibba* is shown in a barplot grouping each 100-nt in size (X axis) and by percentage of the total (Y axis). **(b)** Structure of lncRNAs in selected plants (Dataset S1). The box plot groups the proportion of exon number (X axis) by percentage (Y axis) among lncRNAs identified from various plant genomes. **(c)** Genomic annotation of *U. gibba* lncRNAs. The proportion is from 4,295 putative lncRNAs loci identified. Red represents the proportion of lncRNAs mapped to body gene regions; blue shows lncRNAs located in intergenic regions; green indicates those that overlap gene body and intergenic regions. **(d)** Coding and noncoding transcripts expression. The violin plot represents the total of mRNAs, lncRNAs and lincRNAs identified in this study with correspondent log2(tpm).

To visualize the distribution lincRNAs *loci*, reads were mapped onto the 18 largest contigs of the *U. gibba* genome (>1Mb), which included 4 putatively entire chromosomes. *loci* encoding lncRNAs were distributed across the entire genome with prevalence in high gene density regions, and low frequency in pericentromeric regions (Fig. 2), which can be more clearly seen in the four complete chromosomes (Fig. 2; dark grey color). lncRNAs density was similar in 17 of the largest contigs of the *U. gibba* genome (Lan *et al.*, 2017), except in Unitig_8 (6.8 Mb) that has considerably lower density of lncRNA *loci* (Fig. 2).

**Fig. 2.**
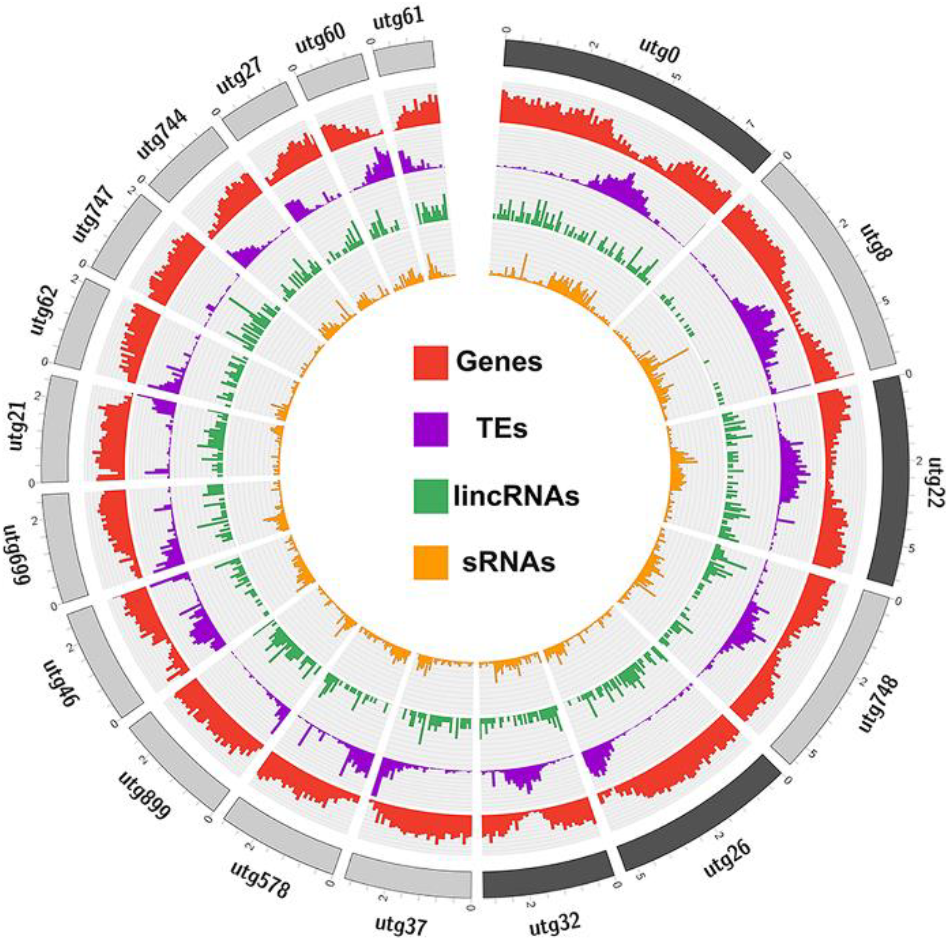
Noncoding RNA landscape in *Utricularia gibba*. Representation of the *U. gibba* genome in a circus plot where the outside blocks represent complete chromosomes (dark gray) and Unitigs or contigs (gray) larger than 1Mb in size. The density plots were calculated in 10Kb windows. Red histogram shows the gene density. The Purple histogram displays transposable element density. lincRNA density is represented in the green histogram, and small RNA density for sRNAs 20-nt to 24-nt in size are shown in orange.

### *U. gibba* has an atypical abundance of 24-nt sRNAs compared to other angiosperms

To further characterize noncoding RNA diversity in *U. gibba*, we carried out small RNA-Seq analysis of RNAs extracted from green tissue and traps from plants grown under the same conditions described for lncRNA identification. We obtained a total of 23.6 million mapped reads, of which 19.9 million had lengths between 20 to 25-nt, which were selected for further analyses (Dataset S1). Upon mapping the reads onto the *U. gibba* genome, we found that *loci* for 20 to 25-nt sRNAs are mainly located at pericentromeric regions, where TE density is higher, but with some peaks in regions of high gene content (Fig. 2). Similar results have been published previously for other plant species (Morin *et al.*, 2008).

An interesting finding was that 52.8% of sRNA sequencing reads corresponded to 21-nt sRNAs and only 14.4% to 24-nt sRNAs (Fig. S4; Dataset S1). This contrasts with sRNA size distribution for other angiosperms, for which the most abundant sRNA class is 24-nt (Chen *et al.*, 2010; Montes *et al.*, 2014). To confirm that the sRNA size distribution in *U. gibba* differs from that of other angiosperms, we performed an analysis of sRNA size abundance in representative plants from different clades for which sRNAs have been characterized (Dataset S1). In total, we analyzed sRNA datasets for 30 plant species, of which 21 were angiosperms and 9 were representative plants outside the angiosperms. Our results confirm that in both monocot and eudicot species, except for *U. gibba*, the most abundant sRNA class is 24-nt (Fig. 3a). For the case of green algae, the most prevalent sRNAs classes are of 21 to 23-nt, whereas in *Volvox carteri* the most abundant sRNAs are of 21-nt and 22-nt, in *Chara coraline* the most abundant are 22-nt and 23-nt (about 30% of each size), and for *Chlamydomonas reinhardtii* the 21-nt class predominates (Fig. 3a). In early-branching land plants (*Marchantia polymorpha, Physcomitrella patens*, and *Marsilea quadrifolia*) and gymnosperms (*Picea abies*, *Gynkgo biloba*) the most abundant size class of sRNAs is 21-nt, except for *Cycas rumphii*, which has similar amounts of 21 and 24-nt sRNAs (Fig. 3a).

**Fig. 3.**
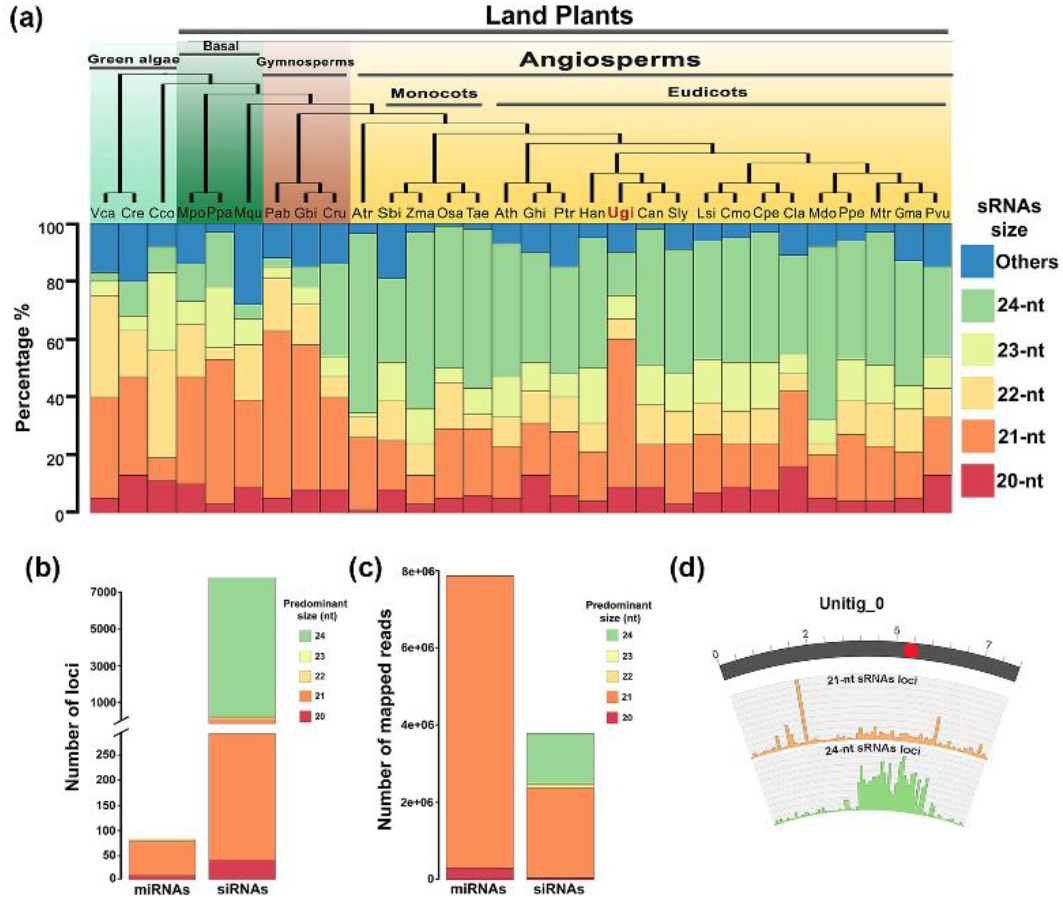
Annotation of sRNAs in *U. gibba* and distribution of small RNA size abundance in plants. **(a)** Top, a phylogenetic tree of plants with representative species for green algae, early branching land plants, gymnosperms, and angiosperms. Bottom, a stacked plot of small RNA abundances (20-nt to 24-nt in size) from various small RNA-seq studies (ataset S1). **(b)** Total locus predictions from ShortStack for miRNAs and siRNAs at different sizes. **(c)** Numbers of reads mapped per loci at different sizes from ShortStack. **(d)** The largest complete chromosome (Unitig_0) representation of gene density, TE density, 21-nt small RNAs loci density distribution and 24-nt small RNAs loci distribution. *Vca: Volvox carteri; Cre: Chlamydomonas reinhardtii; Cco: Chara coraline; Mpo: Marchantia polymorpha; Ppa: Physcomitrella patens; Mqu: Marsilea quadrifolia; Pab: Picea abies; Ginkgo biloba; Cru: Cycas rumphii; Atr: Amborella trichopoda; Sbi: Sorghum bicolor; Zma: Zea mays; Osa: Oryza sativa; Tae: Triticum aestivum; Ath: Arabidopsis thaliana; Ghi: Gossypium hirsutum; Ptr: Populus trichocarpa; Han: Helianthus annuus; Ugi: Utricularia gibba; Can: Capsicum annuum; Sly: Solanum lycopersicum; Lsi: Lagnaria siceraria; Cmo: Cucurbita moschata; Cpe: Cucurbita pepo; Cla: Citrullus lanatus; Mdo: Malus domestica; Ppe: Prunus persica; Mtr: Medicago truncatula; Gma: Glycine max; Pvu: Phaseolus vulgaris*.

### A large number of 24-nucleotide small RNA *loci* produce a small proportion of sRNA reads and are associated preferentially with intergenic regions in *U. gibba*

In general, sRNA sequence distribution in angiosperms is characterized by a major 24-nt peak containing primarily unique reads, and a 21-nt peak comprising many redundant reads (Lelandais-Brière *et al.*, 2010). As expected this is also true in angiosperm species such as Arabidopsis (Wang *et al.*, 2011), tomato (Gao *et al.*, 2015) and rice (Jeong *et al.*, 2011). For the three samples we sequenced, 21-nt sRNAs had higher redundancy (up to 97% of the reads are redundant) in comparison with 24-nt sRNAs, of which 30% were unique and 70% redundant (Dataset S1). To better classify sRNA *loci*, we performed an analysis with ShortStack V2.0 (Axtell, 2013) to identify *in silico* which *DICER*-like (*DCL*) genes are involved in the biogenesis of miRNAs or siRNAs. ShortStack analysis identified 7478 siRNA *loci* and only 80 miRNA *loci* (Fig. 3b; Dataset S3), which produce nearly 1.5 million and 8 million mapped reads respectively, suggesting that there is large number of low abundant siRNA *loci* and few miRNAs that are present in high levels (Fig. 3c). Of the 80 miRNA *loci* identified, 78 already were annotated in the miRBase V22.0 catalog of eukaryotic miRNAs, and 2 represent putative Utricularia specific miRNAs (Dataset S3; Fig. S5). miRNA *loci* grouped into 17 families, of which the miR166, miR156, miR159, miR319 and miR858 families made up 94% of the miRNA reads (Fig. S6). These miRNA families are conserved in most plant species and produce high levels of mature miRNAs (Montes *et al.*, 2014). sRNA *loci* annotation reveals that 80% of 24-nt sRNA *loci* are preferentially located at IGRs, while the sRNA *loci* of 20-nt to 22-nt were located in similar proportions at genic and intergenic regions, while the 23-nt sRNA class, 57% of *loci* are located at IGRs and 31% in genic regions (Fig. S7).The annotation is consistent with the distribution of sRNAs at the genome scale and can be clearly seen in unitig_0, in which 21-nt sRNA *loci* are distributed across the chromosome with significant peaks in regions with high gene density and 24-nt sRNA *loci* were found to be predominantly located in regions with high TE density, presumably pericentromeric regions (Fig. 3d).

### sRNA biogenesis and the RdDM pathway in *U. gibba*

The unusual proportions of 24-nt and 21-nt sRNAs observed in *U. gibba* suggest that its sRNA production machinery could differ somehow from those of other angiosperms. To explore this possibility, we focused on the presence in the *U. gibba* genome of genes involved in canonical miRNA biogenesis, genes that are part of the subunits of RNA Pol-IV and RNA Pol-V, homologs of AGO, DCL, RDR, and other key genes involved in siRNA biogenesis and *de novo* DNA methylation. We searched based on sequence homology (transcript and protein), protein domain conservation, synteny, and through phylogenetic analysis. Furthermore, we performed a comparison with representative plant species (both angiosperm and non-angiosperm) for which key RdDM pathway genes (Ma *et al.*, 2015) and RNA polymerase compositions were previously reported (Huang *et al.*, 2015).

We found that key genes involved in canonical miRNA biogenesis such as *DICER LIKE 1 (DCL1)*, *SERRATE* (*SE*), *HYPONASTIC LEAVES 1* (*HYL1*), *HASTY 1* (*HST1*) and *ARGONAUTE 1* (*AGO1*) are conserved in *U. gibba* (Table S2). Pol-IV and Pol-V are crucial in the RdDM pathway and are constituted by diverse DNA-DIRECTED RNA POLYMERASES IV AND V (NRPD/NRPE) subunits. Pol-IV and Pol-V differ from RNA polymerase II in their second, fourth, fifth, and seventh subunits (NRPD2, NRPD4, NRPE5, NRPD7, respectively). The *U. gibba* genome encodes *NRPD1, NRPE1 and NRPD2* genes (Fig. S8), which is consistent with the presence of siRNAs and their strong conservation in the land plant lineage. We also identified NRPD7 in the *U. gibba* genome (Fig. S8), a subunit previously identified in green eukaryotes except in the algae Chlamydomonas, and NRPD5, found in gymnosperms and angiosperms but not in algae and ferns (Huang *et al.*, 2015; You *et al.*, 2017). Only after an extensive search did we find evidence for an NRPD4 ortholog in *U. gibba*. Although this gene is classified as an orphan in Plaza 4.0 (HOM04D168668) for *Arabidopsis thaliana*, our analysis showed that this categorization is likely related to the *Arabidopsis thaliana* homolog being extremely divergent; even the *A. lyrata* ortholog was readily placed into a clear NRPD4 gene family (Fig. S9). Interestingly, this subunit has been reported only in angiosperms and not in gymnosperms, early land plants or algae (Huang *et al.*, 2015; You *et al.*, 2017).

The AGO protein family in Arabidopsis has been subdivided into 4 clades: AGO2/3/7, AGO4/6/8/9, AGO1/10 and AGO5. The number of family members of this protein family ranges from 2 AGO proteins in *C. reinhardtii* to 10 in Arabidopsis and 20 in *Z. mays* (Rodríguez-Leal *et al.*, 2016). We searched in the Phylome database v4 (Huerta-Cepas *et al.*, 2013) for possible homologous genes in *U. gibba* and found evidence for at least one AGO for each clade, with the exception of AGO5 for which no homologs were found (Fig. S10). Two genes were found for AGO clade 2/3/7 (Fig. S11), two genes represented the AGO 4/6/8/9 clade (Fig. S12), and 4 homologs grouped in the AGO 1/10 clade (Fig. S13). Additionally, we performed our own exhaustive phylogenetic analysis with many plant genomes to assign each *U. gibba AGO* gene with more certainty into specific clades. We were able to identify the same 8 AGOs described above, wherein the 2 homologs of clade 2/3/7 apparently are AGO7 copies, the two 4/6/8/9 copies appear closer to AGO4 than AGO6. AGO8/9 are sister genes only in Brassicaceae. There is one *U. gibba* AGO10 and three homologs for AGO1 in the AGO1/10 clade (Fig. S14).

In seed plants there are three *RDR* ortholog genes; two are conserved in all land plants, *RDR1* and *RDR6,* and *RDR2* required for production of Pol-IV-siRNAs that is specific to seed plants. Aside these RDRs, three additional members of this protein family, RDR3, RDR4, and RDR5, are present in Arabidopsis and other plants (Willmann *et al.*, 2011; Hunter *et al.*, 2016; Qin *et al.*, 2018). We found RDR6 (two copies), RDR1 and RDR2, but no evidence for the presence of RDR3/4/5 in the genome of *U. gibba* (Fig. S15a). In angiosperms the DCL family has 4 members: DCL1, DCL2, DCL3, and DCL4 (Mukherjee *et al.*, 2013); however, in lycophytes and ferns there is only evidence for the presence of DCL1, DCL3 and DCL4 (You *et al.*, 2017). Phylogenetic analysis of the DCL family permitted the identification in *U. gibba* of 4 DCL proteins (DCL1, DCL2, DCL3, DCL4), suggesting that has a DCL repertoire similar to other angiosperms (Fig. S56b). Globally, were able to assign *U. gibba* homologs for the remaining key genes in the canonical RdDM pathway (Dataset S1). However, although UgDCL3 phylogenetically groups very closely to tomato and Mimulus DCL3, UgDCL3 is missing its N-terminal region, where the conserved DEAD/DEAH, Helicase and Dicer dimerization domains are located (Fig. 4a; Fig. S16). The absence of about 600 amino acids of the N-terminal region in UgDCL3 is evident in a multiple sequence alignment of DCL3 with other angiosperms (Fig. 4b; Fig. S17). To confirm that the incomplete DCL3 does not represents mistake in the assemble/annotation of the *U. gibba* genome, we searched for the presence of DCL3 transcripts in the different RNA-Seq libraries and we confirm with a 5’ Rapid Amplification of cDNA Ends (5’RACE) that in both cases the sequence of transcripts and 5’ RACE sequence that the *U. gibba* DCL3 gene is missing the DEAD/DEAH domain (Fig. S18).

**Fig. 4.**
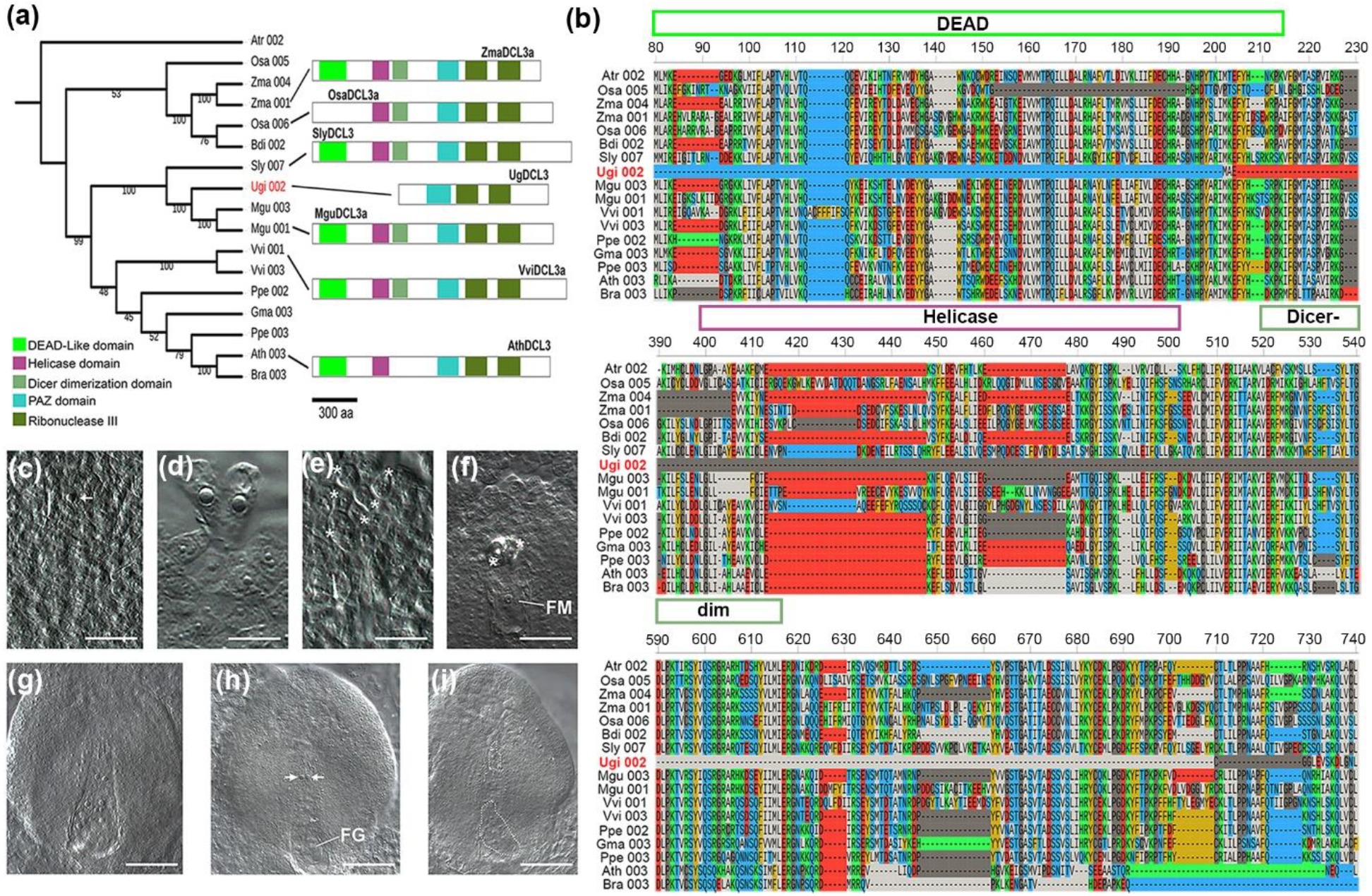
Truncated DCL3 and female gametophyte development in the ovule of *U. gibba.* **(a)** Phylogenetic analysis of DCL3 in some angiosperms including *U. gibba*. A typical DCL3 protein has the DEAD, Helicase, Dicer dimerization, PAZ, and two Ribonuclease III domains, while UgDCL3 only contains the PAZ and Ribonuclease III domains. **(b)** Multiple sequence alignment of DCL3 proteins in blocks of 150 aa showing the missing domains in UgDCL3. **(c)** Developing ovules showing a single pre-meiotic precursor corresponding to the MMC. **(d)** Twin pre-meiotic precursors prior to megasporogenesis. **(e)** Six differentiated cells (asterisk) corresponding to gametic precursors prior to megasporogenesis. **(f)** Functional megaspore (FM) and two degenerated megaspores (asterisk) following megasporogenesis. **(g)** Developing ovule showing a single 8-nuclear female gametophyte. **(h)** Developing ovule showing a female gametophyte (FG) and two ectopic gametic precursors (arrows) at the chalazal region. **(i)** Developing ovule showing two independent female gametophytes (dashed). Scale bars: 12.5 μm in **(c)** to **(f)**; and 20 μm in **(g)** and **(i)**.

### Female gametogenesis in *U. gibba* is reminiscent of Arabidopsis mutants affected in the RdDM pathway

Mutations affecting most of the genes involved in the RdDM pathway have no obvious phenotype during the vegetative development of plants, but show defects in female gametogenesis, including the differentiation of supernumerary gametic precursors that often give rise to ectopic female gametophytes within the ovule (Olmedo-Monfil *et al.*, 2010; Hernández-Lagana *et al.*, 2016). To determine if the truncation of DCL3 and the unusual distribution of sRNAs could be related with altered female gametogenesis in *U. gibba* as has been reported for Arabidopsis mutants affected in the RdDM pathway, we analyzed female gametogenesis in whole-mounted developing ovules. No descriptions of ovule development have been previously reported for *U. gibba*. Our results are described and illustrated in Fig. 4c-i and in Table S3.

As for other species of *Utricularia* (Jhori BM, Ambegaokar KB, 1992; Płachno & Swiątek, 2012), the ovule of *U. gibba* is unitegmic, with a funiculus forming a raphe and merging into a voluminous placenta. The formation of differentiated gametes occurs after the formation of meiotically derived megaspores (megasporogenesis). Subsequent rounds of mitotic divisions give rise to the female gametophyte (megagametogenesis). Megasporogenesis occurs in ovules within ovaries having a diameter of 0.3 to 0.5 mm. Whereas 51.4% (n=142) of pre-meiotic ovules showed a single megaspore mother cell (MMC; Fig.4c), 42.9% showed from two to six differentiated cells resembling the MMC (Fig. 4d and e). In 5.7% of the ovules examined we could not identify a pre-meiotic precursor. Ovules contained in 0.5-1 mm ovaries had already undergone meiosis and often show a chalazal functional megaspore (FM; Fig. 4f) within which mitotic divisions will give rise to an 8-nucleated syncytium (Fig. 4g) that will cellularize before differentiating into a mature female gametophyte (FG). Although the degeneration of the FG prior to the end of megagametogenesis is not uncommon (13.1%; n=76), in most cases (40.7%; n=76) the ovule contains a FG in which the micropylar region containing the egg apparatus expands outside the integument and grows within the placenta (Fig. 4g and 4h). Interestingly, 22.4% of ovules examined showed supernumerary gametic cells in the chalazal region, independent of the developing FG (Fig. 4h), and containing two independently developing FGs (Fig. 4i), suggesting that supernumerary gametic precursors can give rise to female gametophytes that may or may not originate from a meiotically derived cell. The presence of supernumerary gametic precursor cells and ectopic female gametophytes in *U. gibba* is reminiscent of phenotypes found in Arabidopsis mutants *dicer-like3* (*dcl3*), *argonaute4* (*ago4*), *argonaute9* (*ago9*), *rna-dependent rna polymerase6* (*rdr6*), and *nrpd1a*, all affected in key components of the RdDM pathway.

### *U. gibba* has a reduced levels of DNA methylation

In plants, 24-nt siRNAs from repetitive DNA and TEs that are loaded by AGO4/6 trigger DNA methylation, which results in histone modifications such as the H3K9me2 (Lippman *et al.*, 2004; Xu *et al.*, 2013). Because of the unusual distribution of 24-nt sRNAs and the low proportion of TEs and other repetitive DNA in *U. gibba*, we decided to explore preliminarily the global DNA methylation patterns in this carnivorous plant using long-read DNA sequencing data with the technology of SMRT-Seq, recently reported (Lan *et al.*, 2017). This technology measures each base addition as an interpulse duration (IPD) or retention time ratio. The IPD will depend on whether the new base is incorporated by pairing to a modified or non-modified base in the template and the nature of the modification (Flusberg *et al.*, 2010). Therefore, analysis of the IPDs during SMRT-Seq can allow the identification of m5C in the DNA template without the need for commonly used bisulfite DNA chemical conversion methods. Since there are no previous reports of using PacBio sequencing data to determine m5C global methylation levels in plants, we tested whether the IPDs could be used for m5C identification in *Arabidopsis thaliana*. With this aim we analyzed the IPDs from 40 cells of SMRT-Seq data obtained from Arabidopsis plants grown under normal conditions (Methods S1) and compared with BS-Seq data previously reported for Arabidopsis plants grown similarly (Yong-Villalobos *et al.*, 2015). In global terms, the percentage of cytosines covered in the SMRT cells was 95.51%, while for BS-seq it was 99.95% (Table S4). Mean coverage for this base was 46.9X for the SMRT data and 15X for BS-Seq (Methods S2-S3). The total number of modified cytosines detected in the SMRT data was 9.21% (3,906,986), which is higher than the 7.71% of 5mC determined by BS-Seq (Table S4). To obtain greater confidence in obtaining *bona fide* m5C from SMRT data, we selected only those with IPD equal or above 1.79, resulting in 5.08% m5C (2,047,109), which was slightly lower than that of BS-Seq. To confirm that the global methylation in *U. gibba* determined by SMRT data indeed represent what can be obtained by the BS-Seq, we carried out correlation analysis of methylation for each chromosome. We found a high global Pearson’s correlation with r=0.9042 (P-value P ≤ 2.26 × 10^−35^) between methods, which varied from r=0.85 for chromosome 1 to r=0.95 for chromosome 3 (Fig. S19a and Table S5). The high correlation obtained between the two methods suggests that SMRT data should allow to obtain representative image of the methylation landscape of plant genomes. The global methylation context for SMRT data was calculated to be 42.8% CG, 20.9% CHG, and 35.9% CHH, values that are within the ranges previously reported for Arabidopsis BS-Seq data: 39-41.5 for CG, 17-24% CHG and 36-41% to CHH (Fig. S19b) (Cokus *et al.*, 2008; Dowen *et al.*, 2012; Yong-Villalobos *et al.*, 2015). When BS-Seq and SMRT-Seq global methylation patterns were graphically compared at the chromosome level, the distributions of methylated regions were similar and the characteristic peaks of dense methylation at centromeric regions were consistent between both methylomes (Fig. S19c).

Since SMRT-Seq analysis showed a similar pattern and distribution of m5C in the Arabidopsis genome, we used the same strategy to generate a preliminary methylome estimation in *U. gibba*. The genome-wide depth with SMRT sequencing of *U. gibba* was ~ 70X and the mean coverage for all bases was 34X. We identified 1,590,729 putatively methylated cytosines and 1,088,032 high-confidence m5Cs (IPD >= 1.7), which represents 3.88% and 2.69% of the Cs in the *U. gibba* genome, respectively. DNA methylation corresponding to all methylated cytosines in high confidence m5Cs for each context was scored: 37.30% CG methylation, 22.22% in CHG context, and 40.45% in CHH (Fig. S20).

Because of the unusually low level of m5C detected by SMRT-Seq analysis, we performed BS-Seq for two replicates of the whole-plant of *U. gibba*. The bisulfite conversion rates in both replicates were greater than 99.85% and the mean coverage for base was 26X. After sequencing, the clean reads were mapped against the reference genome obtaining around 85% of mapping rate. During the base calling we found a total of 1,281,545 and 1,302,422 high confidence m5C for each replicate. In percentages of total cytosines in the genome, the global methylation ranged from 2.92%-3.17% from and high-confidence m5C. The distribution of m5Cs at the chromosome level shows peaks near centromeric regions for both replicates of BS-Seq and for SMRT-Seq (Fig. 5a) with very similar distributions, similar to that reported in other plant genomes but with one of the lowest global methylation rates reported for a land plant (Niederhuth *et al.*, 2016; Vidalis *et al.*, 2016). Of the total of m5Cs identified, 38.8%, corresponds to CG context, 27.3% to CHG and 33.86% to CHH (Fig. 5b). The percentage of m5C present in the CG, CHG and CHH context respect to the total number of these contexts in the genome were 12% for CG, 8% for CHG and 2% for CHH (Fig. 5c). These percentage of methylation are lower as compared to other angiosperms including *A. thaliana* (Takuno *et al.*, 2016). To explore gene body methylation (GbM) in *U. gibba*, we assessed the methylation patterns for gene bodies and upstream/downstream (0.8 Kb) regions for both genes and TEs. In genic regions we found lower DNA methylation density levels in upstream/downstream regions than in gene bodies, with major peaks located near start/stop codon sites with CG methylation being the highest in the gene body region. (Fig 5d). These results correlates with GbM densities previously reported for other plants (Takuno *et al.*, 2016; Bewick & Schmitz, 2017). The analysis of TE body methylation showed lower methylation levels near upstream and downstream regions such as in GbM. However, TEs had higher methylation density in CHH rather than CG contexts (Fig. 5e), which could explain TE silencing in *U. gibba*.

**Fig. 5.**
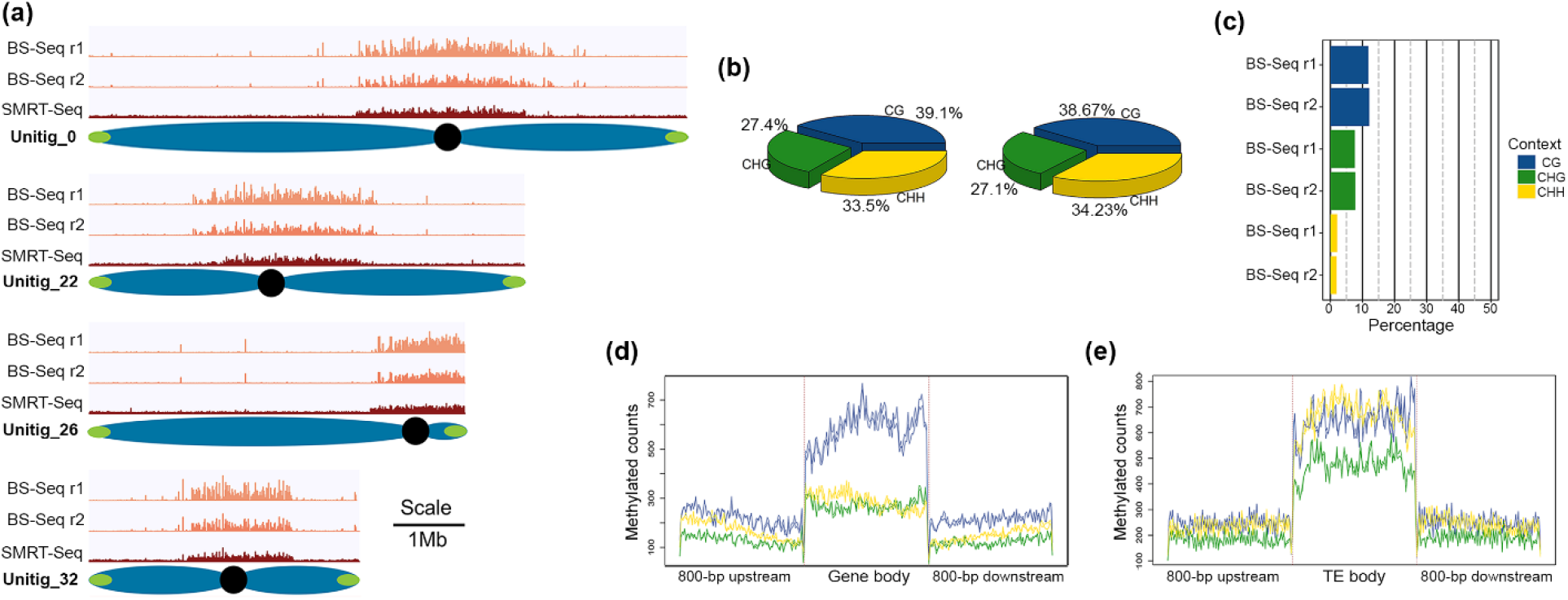
Genome-wide methylation in *Utricularia gibba*. **(a)** Distribution of methylation (m5C) representation only in the complete chromosomes in *U. gibba* for location of pericentromeric regions. Here, the blue light red shows the methylation density in BS-Seq replicates and the red histogram represent the methylation distribution in SMRT-Seq with normalized data (Scale 1Mb). **(b)** Methylation context at the genome level for two replicates. CG methylation in blue, CHG methylation in green, and CHH methylation in yellow. **(c)** Methylation levels for CG, CHG and CHH context. **(d)** Gene body methylation and TE body methylation representation in all contexts (CG, CHG and CHH) 800bp upstream and downstream. **(e)** TE body methylation and TE body methylation representation in all contexts (CG, CHG and CHH) 800bp upstream and downstream.

## Discussion

### Genome rearrangements after WGD in *U. gibba* and their role in ncRNA content

The causes and mechanisms of WGD events and genome fractionation processes that lead to genome expansion and contraction in plants, are still poorly understood. Moreover, the consequences of these processes on the repertoire of non-coding RNAs and epigenetic processes remain obscure. *U. gibba*, a carnivorous plant with an unusually small but dynamic genome that has experienced two relatively recent WGD events, represents a highly illustrative model to study the processes of genome contraction and its consequences on the diversity of ncRNAs and epigenetic processes. Our analysis of ncRNAs provides insights into possible reasons why this carnivorous plant has an unusual siRNAs distribution. One of our interesting observations is that in contrast to other angiosperms, such as Arabidopsis, maize, and tomato, for which lncRNA *loci* are preferentially located in centromeric regions, in *U. gibba* lncRNA *loci* are well distributed across the genome but with a lower density in centromeric regions (Fig. 2). Centromeres are genomic sites of spindle attachment essential for ensuring proper chromosome segregation during cell division. Despite their recognized functional importance, centromeres are not well defined at the sequence level in eukaryotic genomes except for some small fungal genomes (Buscaino *et al.*, 2010). In general, centromeres are accepted to be composed of high-copy tandem satellite repeats and/or the presence of centromeric chromoviruses, a lineage of Ty3/gypsy retrotransposons. Also, these sequences may be merely parasitic and tend to accumulate in recombination-poor centromeric regions to escape negative selection (Gao *et al.*, 2008). The low density of lncRNAs observed in the putative centromeric regions of the *U. gibba* genome could be related to the absence of both satellite repeats and paucity of centromeric chromoviruses in these regions (Lan *et al.*, 2017). Centromeres without long arrays of satellite DNA have been referred to as evolutionarily new centromeres (Piras *et al.*, 2010). It is possible that after the last WGD event in *U. gibba*, genome fractionation resulted in the formation of neo-centromeres targeted by few lncRNAs. These findings are consistent with the notion that epigenetic marks, in the form of stable, self-propagating chromatin states, rather than sequence specific structural features, define a functional centromere.

### A truncated *DCL3* gene correlates with the distribution of sRNAs and impact the developmental biology of *U. gibba*

TE proliferation shapes genome sizes among eukaryotes. In plants, one of the mechanisms to control this expansion is through TE silencing via 24-nt siRNAs (Lisch, 2008; Tenaillon *et al.*, 2010). In the siRNA biosynthesis pathways, at least three RNA-dependent RNA polymerases (RDR1, RDR2, and RDR6) are functional in plants (Willmann *et al.*, 2011; Hunter *et al.*, 2016). RDR2 works with DCL3 to form chromatin associated siRNAs (mainly 24-nt) that are involved in sRNA mediated DNA methylation (Matzke & Mosher, 2014). As expected from its known role in the biogenesis of 24-nt small RNAs, Arabidopsis and maize mutants are defective in RDR2 or DCL3 show a severe reduction in the production of the 24-nt size siRNA class, whereas the 21-nt class (miRNA) is not affected and indeed becomes overrepresented (Kasschau *et al.*, 2007; Nobuta *et al.*, 2008). In rice, DCL3 mutations also affect the production of 24-nt siRNAs associated with TEs (Wei *et al.*, 2014). Here, we report that *U. gibba* has an unusually a sRNA size distribution with low content of 24-nt sRNAs, more similar to early-branching land plants and gymnosperms than to other angiosperms (Fig. 3a). The accumulation of 24-nt siRNAs in *U. gibba* is similar to Arabidopsis, rice and tomato *dcl3* mutants (Kasschau *et al.*, 2007; Kravchik *et al.*, 2014; Wei *et al.*, 2014), which correlates well with the presence in *U. gibba* of a truncated most likely inactive form of DCL3 (Fig. 4a,b). The loss of a fully functional DCL3 in *U. gibba* may be responsible for the reduction in the 24-nt class of siRNAs, as well as the reduced level of DNA methylation in its genome. It is possible that the loss of a fully functional DCL3 in *U. gibba* might have originally reflected lower selective pressure to silence fewer TEs after genome size reduction. Furthermore, we report the first quantitative analysis of female gametogenesis in a member of the *Utricularia* genus showing developmental ovule defects that have not been reported previously for *Utricularia* species (Płachno, 2011; Płachno & Swiątek, 2012).

Since the truncated DCL3 is structurally preserved by purifying selection, some function, perhaps regulatory, likely remains. Animal Dicer and plant DCL proteins dimerize at their RNase III domains (Zhang *et al.*, 2004; Mickiewicz *et al.*, 2017). It is possible that the truncated DCL3 acts as an interfering subunit in the DCL protein complex to generate a dominant negative phenotype (Veitia *et al.*, 2018). As hypothesis, this dominant negative effect could suppress the production of the 24-nt class of siRNAs accompanied by defects that include the absence of a female gametophyte that leads to partial female sterility, or the differentiation of female ectopic precursors that give rise to supernumerary gametophytes. It would be interesting to further study the biological implications of the DCL truncation in *U. gibba*, as it could represent an initial step in the evolution of apomixis and further developmental and genetic studies are required to determine if the latter could eventually result in variable frequencies of unreduced gamete formation, polyploidization or apomixis.

### SMRT-Seq as an alternative to decipher plant methylomes

All plant methylomes sequenced to date have been generated by BS-Seq (Flusberg *et al.*, 2010). In recent years, however, new techniques for sequencing has generated the possibility of direct DNA base modification detection without any chemical conversion (Flusberg *et al.*, 2010). Recently, genome-wide DNA methylation on N-6 adenine (m6A) using SMRT-sequencing technology was reported for some animals and plants (Greer *et al.*, 2015; Wu *et al.*, 2016; Liang *et al.*, 2018). Our results of *U. gibba* BS-Seq support the use of SMRT sequencing technology to determine m5Cs on whole genomes. At least in qualitative terms, SMRT technology produces whole-genome m5C data similar in the percentages of methylated cytosines, the proportions of methylation context (CG, CHG and CHH), genome-wide m5C distribution and GbM to those obtained by bisulfite sequencing. BS-Seq has so far been the standard for methylome determination and has been quite useful to derive insights into the roles of DNA methylation on gene expression and chromatin remodeling. However, because BS-Seq is based on short-length sequencing reads that provide a statistical probability that a cytosine is methylated, it is difficult, if not impossible to determine which m5C modifications need to be present in the same molecule to influence gene expression and/or chromatin remodeling. Although further analyses will be required to ascertain the extent to which SMRT technology can generate robust quantitative data on m5C DNA modifications, the fact that methylation patterns can be established linearly for long-reads could serve to establish which m5C modifications are truly necessary to influence gene expression or chromatin modifications.

### Lower DNA methylation levels in *U. gibba* in comparison with other plant genomes

Global DNA methylation of *U. gibba* ranges from 2.69 to 4.09%, lower than that reported for other plant species, which range between 5% in *Theobroma cacao* to 43% in *Beta vulgaris* (Vidalis *et al.*, 2016). Our results suggest that the reduced level of 24-nt siRNAs in *U. gibba* could be due to the presence of a truncated DCL3 and/or the absence of AGO6, in turn impacting DNA methylation. In spite of having lower levels of DNA methylation, CG methylation in *U. gibba* is similar to that of Arabidopsis in its preferential localization in gene bodies and heterochromatic regions, and opposite to that found in Selaginella, where CG methylation is located mainly outside gene body regions (Zemach *et al.*, 2010). Interestingly, the proportion of methylated genes with GbM was similar than in Arabidopsis (30%) and in early land plant (Bewick & Schmitz, 2017). We suggest that GbM in *U. gibba* could serve as a mark for DNA sequences that need to be maintained after drastic genome reorganization, such as broad deletional DNA loss following its successive WGD events.

## Acknowledgments

We thank G. Alejo-Jacuinde, B. Pérez-Sánchez and Q. Ortiz-Vasquez for help collecting plants and flowering specimens. Special thanks to R. Purbojati and S. Schuster for providing us the PacBio raw sequencing data. We thank J. Cervantes-Luevano for informatic support, Luis Delaye for evolutionary analysis advice and Koribinian Schneeberger for provided us syntenic dataset generating by SyRI. Authors acknowledge Dr. T. Markow for critical review of this manuscript. S.A.C.-P., I.A.-M. and F.R.-R are the recipients of a graduate scholarship from Conacyt. This work was supported in part by a Senior Scholar grant from the Howard Hughes Medical Institute and a Basic Science grant from CONACytT Mexico to L.H.-E.

## Author contributions

S.A.C.-P. and L.H.-E. designed research; S.A.C-P., L.Y.-V, N.M.V.F.-Z, A.O.-A., I.A.-M., and F.R.-R. performed research; L.H.-E., VAA and J.P.V.-C. contributed new reagents/analytic tools; S.A.C.-P., L.Y.-V., N.M.V.F.-Z., T.L, O.M., J.P.V.C., V.A.A., and L.H.-E. analyzed data; and S.A.C.-P., VAA and L.H.-E. wrote the paper with inputs from all authors.

Note: Supplementary Materials and datasets accession number will be available after acceptance.

## Notes

### Competing Interest Statement

The authors have declared no competing interest.

